# The effect of hunger and state preferences on the neural processing of food images

**DOI:** 10.1101/2025.09.09.674354

**Authors:** Denise Moerel, Cecilia Chenh, Sophie A. Bowman, Thomas A. Carlson

**Affiliations:** School of Psychology, University of Sydney, Sydney, Australia; The MARCS Institute for Brain, Behaviour and Development, Western Sydney University, Sydney, Australia

## Abstract

Visual information plays a key role in guiding food-related decisions. While previous studies have shown that features such as calories and naturalness are encoded by the brain, upon simply seeing the stimuli, it remains unclear how this encoding is shaped by the observer’s current state. In this study, we explore the effect of 1) hunger state, 2) task relevance, and 3) current individual preference on the processing of visual food information. Participants (N = 23) underwent two EEG sessions: one after fasting overnight and another after eating normally. During each session, participants did two separate tasks, one where the stimuli were task-relevant and one where attention was distracted away. We used multivariate analysis methods to assess the impact of hunger on the representation of food-related features, and to determine the time-course of information related to food flavour, personal appeal, and arousal, across both tasks. Results showed that information about edibility (food versus non-food object), food identity (e.g. hamburger versus pizza), flavour profile, or personal appeal and arousal was not influenced by the hunger manipulation. Flavour was represented regardless of attentive state, whereas personal appeal and arousal information emerged later and were only observed when the food was task relevant. We found that food appeal and arousal encoding were more closely aligned with behavioural ratings within rather than between sessions, suggesting the nature of the encoding was driven by current state. The study provides insights into how personal preferences and physiological states influence the representation of food information in the brain.

## 1. INTRODUCTION

Humans rely on smell, taste, and vision to perceive, categorise, and choose foods. Among the senses, vision is unique in that it allows us to gather information about foods at a distance. Whether the food is sitting on a plate in front of us or 20m away on a billboard, the brain’s visual system enables us to rapidly gather information to inform our decisions. Evidence from functional Magnetic Resonance Imaging (fMRI) suggests that the brain uses similar mechanisms to encode food cues through vision and taste (Avery et al., 2021; Simmons et al., 2005). In addition, recent fMRI studies have provided evidence that food stimuli may drive the coarse scale organisation of the ventral visual stream (Jain et al., 2022; Khosla et al., 2022). Recent studies have also provided insight into how different features of food are encoded in the brain. For example, multiple studies have shown the brain distinguishes between edible and inedible (i.e., food and non-food) objects using fMRI (Avery et al., 2021; Chen et al., 2016; Killgore et al., 2003; Rothemund et al., 2007; Simmons et al., 2005) and magnetoencephalography (MEG) or electroencephalography (EEG) (Moerel, Psihoyos, et al., 2024; Stingl et al., 2010; Tsourides et al., 2016). An fMRI study further showed that the visual cortex carried information about the taste of the food (sweet, salty, or sour) when viewing food images (Avery et al., 2021). In addition, studies have shown that the neural response to food stimuli is modulated by the energetic/caloric content of the food (Chae et al., 2025; Killgore et al., 2003; Mengotti et al., 2019; Moerel, Psihoyos, et al., 2024; Toepel et al., 2009), and by whether food is natural or prepared (Coricelli et al., 2019; Moerel, Psihoyos, et al., 2024; Pergola et al., 2017; Vignando et al., 2019). However, it is still unclear how the encoding of visually presented food by the brain is mediated by the state of the observer. In this study, we investigate the effect of 1) hunger state, 2) task relevance on the neural coding of visual food information, and 3) current personal appeal, which could be influenced by both temporary cravings and long-term preferences.

Several studies have found an effect of hunger on the neural response to food stimuli. For example, fMRI studies have shown a greater response to task-relevant pictures of foods when participants had fasted (Charbonnier et al., 2018; Cheng et al., 2007; Führer et al., 2008; Holsen et al., 2005; LaBar et al., 2001; Mohanty et al., 2008). In a meta-analysis of fMRI studies, van der Laan and colleagues (2011) found an influence of hunger on visual food stimulus activation in the right parahippocampal gyrus and amygdala, areas suggested to process reward, and left lateral OFC, suggested to process the expected pleasantness of the food. EEG studies also have found effects of hunger on the activation in response to food stimuli (Ilse et al., 2020; Nijs et al., 2010; Stockburger et al., 2008, 2009; Sultson et al., 2019) and non-food stimuli (Sultson et al., 2019). Importantly, this effect of hunger was found when the food items were task-relevant (Nijs et al., 2010), during passive viewing (Stockburger et al., 2008, 2009), when attention was actively focused away from the food items (Sultson et al., 2019), and even without explicit awareness about the food item (Ilse et al., 2020), suggesting that this effect may occur automatically. However, it is not yet clear whether hunger can affect the representation of information about the food, as previous work used univariate analysis methods to show differences in activation driven by hunger on fMRI data (Charbonnier et al., 2018; Cheng et al., 2007; Führer et al., 2008; Holsen et al., 2005; LaBar et al., 2001; Mohanty et al., 2008) and EEG data (Ilse et al., 2020; Nijs et al., 2010; Stockburger et al., 2008, 2009; Sultson et al., 2019). It is possible that hunger functionally enhances the representation of food stimuli, but univariate effects of hunger could also be mediated by arousal or inhibitory control without the need for enhanced stimulus representations.

Some work has suggested that hunger effects could be mediated by other food-related factors, such as appeal and calorie content. For example, one fMRI study showed that medial and lateral OFC responses were enhanced for high compared to low appeal foods, but only when the participant was hungry (Piech et al., 2009). In addition, several other fMRI studies have found effects of hunger are mediated by the calorie content of the food (Frank et al., 2010; Goldstone et al., 2009; Siep et al., 2009). It is, however, possible that this interaction is mediated by an attentional bias. That is, hunger could increase the salience of food items, especially if they are highly appealing. Although most studies cannot distinguish between the effects of salience and valence, recent fMRI work suggests that overall activation largely reflects salience, while valence information was reflected in the pattern of activation (Pimpini et al., 2022). In addition, findings about the contribution of Body Mass Index (BMI) have been mixed, with fMRI studies comparing controls with obese individuals and binge-eaters showing variable findings (Ziauddeen et al., 2012). Some univariate fMRI work showed that neural responses to food can be affected by BMI (Volkow et al., 2011), whereas multivariate fMRI work showed no effect of BMI on the neural representation of visually presented food (Pimpini et al., 2022).

By investigating whether focused attention is required for the brain to represent different food-related information, we can identify which representations occur as a consequence of simply perceiving the food stimulus, regardless of task relevance. This can provide insight into how this information is processed, as features that are represented in the absence of selective attention may processed automatically. For example, previous work showed the neural signal has information about edibility (food versus non-food), naturalness and the level of transformation, and the perceived caloric content, even when the food was not task-relevant (Moerel, Psihoyos, et al., 2024). In addition, it is possible that attention mediates the effect of hunger on visual information processing. For example, behavioural work has provided evidence for attentional capture of food over non-food words (Mogg et al., 1998; Placanica et al., 2002) and images (Piech et al., 2010) when participants are hungry. Further evidence for a role of attention in the modulation of hunger effects comes from EEG (Ilse et al., 2020; Nijs et al., 2010; Stockburger et al., 2008). Here, studies have shown evidence for an enhanced N2pc component (Ilse et al., 2020), linked to orienting of spatial attention, and P300 component (Nijs et al., 2010), linked to attention and working memory, for food compared to non-food stimuli when participants were hungry. Previous MEG and EEG work has also shown that attention can affect the representation of visual information generally, with stronger and/or more sustained representations of the information that is within the focus of attention compared to information that is not attended (Battistoni et al., 2020; Goddard et al., 2022; Grootswagers et al., 2021; Kaiser et al., 2016; Moerel et al., 2022; Moerel, Rich, et al., 2024). It is therefore possible that hunger state and attention modulate the processing of information about food.

The findings from EEG work investigating whether the brain encodes personal food appeal information have been mixed (Chae et al., 2025; Moerel, Psihoyos, et al., 2024; Schubert et al., 2021). One study used multivariate pattern analysis on EEG data to show there was information about the subjective ‘tastiness’ of the food from approximately 530 ms after stimulus onset (Schubert et al., 2021). Notably, the same result was found when participants made different judgments about the food, although the onset of this information emerged later, at 740 ms after stimulus onset. We recently used multivariate pattern analysis on EEG data to investigate whether more general food preferences across the population were represented and whether the brain represented the level of arousal in addition to the valence (palatability) of the food (Moerel, Psihoyos, et al., 2024). Our study found neither valence nor arousal information could be decoded from EEG recordings for visually presented food stimuli. The studies differed in terms of whether the valence and arousal ratings were personal or from another group of participants, and whether the participants had fasted. Therefore, in this study we obtained personal appeal models and studied the brain in both a fasted and fed state.

The goal of this study was to investigate the effects of 1) hunger state, 2) current personal appeal, and 3) task relevance on the visual processing of information about food. We hypothesised that if differences in neural activation driven by hunger, previously found with EEG (Ilse et al., 2020; Nijs et al., 2010; Stockburger et al., 2008, 2009; Sultson et al., 2019) as well as fMRI (Charbonnier et al., 2018; Cheng et al., 2007; Führer et al., 2008; Holsen et al., 2005; LaBar et al., 2001; Mohanty et al., 2008), reflect a general hunger driven enhancement of visual processing of food items, the hunger state of the individual would enhance the neural encoding of various food-related features; edibility, food identity, flavour, appeal and arousal. In addition, we hypothesised that if food-related features are encoded due to simply perceiving the food stimulus, independent of task demands, then focused attention would not be required for their neural representation. Finally, we hypothesised that if the neural signal reflects highly personal appeal, rather than more general food preferences across the population (Chae et al., 2025; Moerel, Psihoyos, et al., 2024; Schubert et al., 2021), personal appeal and arousal, based on ratings provided by the individual during the EEG session, should be encoded in the neural signal.

To test these hypotheses, participants took part in two different EEG sessions, once after fasting overnight (‘fasted’: 13.00 ± 2.37 h since eaten), and once after eating as normal (‘fed’: 1.53 ± 0.78 h since eaten). We use two tasks to manipulate whether the stimuli were task relevant, and therefore within the focus of attention. Using multivariate decoding methods on EEG data, we studied the effect of hunger on the coding of edibility and food identity information in the brain. In addition, we used Representational Similarity Analysis to determine the time-course of information about food flavour, appeal and arousal, obtained through ratings at the start of the session.

## 2. MATERIALS AND METHODS

### 2.1. Participants

Twenty-four individuals participated in two experiment sessions. One participant was excluded from the analysis as they reported their last food intake as 13 hours prior to the session in the fed condition. Therefore, data were analysed for 23 participants (14 female/9 male) with a mean age of 22.83 years (SD = 8.45, range = 18-62) and a mean BMI of 21.87 points (SD = 2.65, range = 17.3-28). The sample size was determined using a Bayesian stopping rule (Keysers et al., 2020; Wagenmakers et al., 2018). We began with 23 participants, based on prior work (Moerel, Psihoyos, et al., 2024; N = 20). Using a stopping criterion of ≥80% of time points with BF < 1/6 or BF > 6 (Teichmann et al., 2022), we found that 84.78% of points across the main analyses (Figures 4–6) met this threshold and therefore stopped data collection after the initial sample.

All participants reported normal or corrected-to-normal visual acuity, were able to fast, and reported no history of epilepsy, head injury, neurological disorder or eating disorder. No participants were currently on a diet, however, three participants had dietary restrictions (gluten free, pescatarian, and intermittent fasting respectively). Participants completed additional surveys, that were unrelated to this study, about their self-reported health, knowledge about calories, and information about their culture and language. Participants received a total of $100 AUD for their participation in both sessions. The study was approved by the Human Ethics Committee of the University of Sydney, and participants provided written and verbal consent prior to participating in the experiment.

### 2.2. Stimuli and procedure

#### 2.2.1. Hunger manipulation

We used a within-subjects design to test the effect of hunger on the representation of food in the brain. All participants attended two two-hour EEG sessions, one session for the fasted condition, and one session for the fed condition. The sessions were scheduled on two separate days, ranging from 3–14 days apart (mean = 7.13 days, SD = 3.17 days). The order of the sessions was counterbalanced to minimise order effects. Both sessions were scheduled at the same time of day to minimise variations in circadian rhythm (Poggiogalle et al., 2018; Stockburger et al., 2008). In the fasted condition, participants were instructed to fast for at least eight hours prior to participating in the experiment, based on previous work (Führer et al., 2008; LaBar et al., 2001; Piech et al., 2010), but they were allowed to drink water. Participants in the fed condition were instructed to eat as they normally would prior to the session. To ensure the effectiveness of the hunger manipulation, participants completed subjective physiological ratings at the beginning of each session. Before participating in the EEG experiment, participants were instructed to rate their hunger, thirst and fatigue levels on a one-hundred-point sliding scale, using an iPad, with 100 indicating very hungry/ thirsty/ tired. Participants also stated the time of their last full meal and the time of their last food intake (including snacks). Note that one participant provided the same subjective hunger rating across both the fasted and fed condition, although they complied with the food intake instructions. As removing this participant did not change the results, we have included this participant in the sample for completeness.

#### 2.2.2. Stimuli

The visual stimuli consisted of 280 segmented colour objects images (see Figure 1A). Half of these were food images (70 natural foods, 70 transformed foods) and half were non-food images (70 animate objects, 70 inanimate objects). All food images were selected from the Foodcast Research Image Database (FRIDa) (Foroni et al., 2013). The subset of images used was selected to be familiar for Australian participants, and rotten food items were excluded. We controlled for the transformation of the foods by selecting 70 natural and 70 transformed foods. Natural foods are defined as being in their original state (e.g. banana), while transformed foods are foods whose form has been changed by humans in some way, for example, by cooking or preserving (e.g. donut). For the non-food images, 88 were selected from FRIDa (Foroni et al., 2013), and the remaining 52 images were obtained from PNG images.

**Figure 1.**
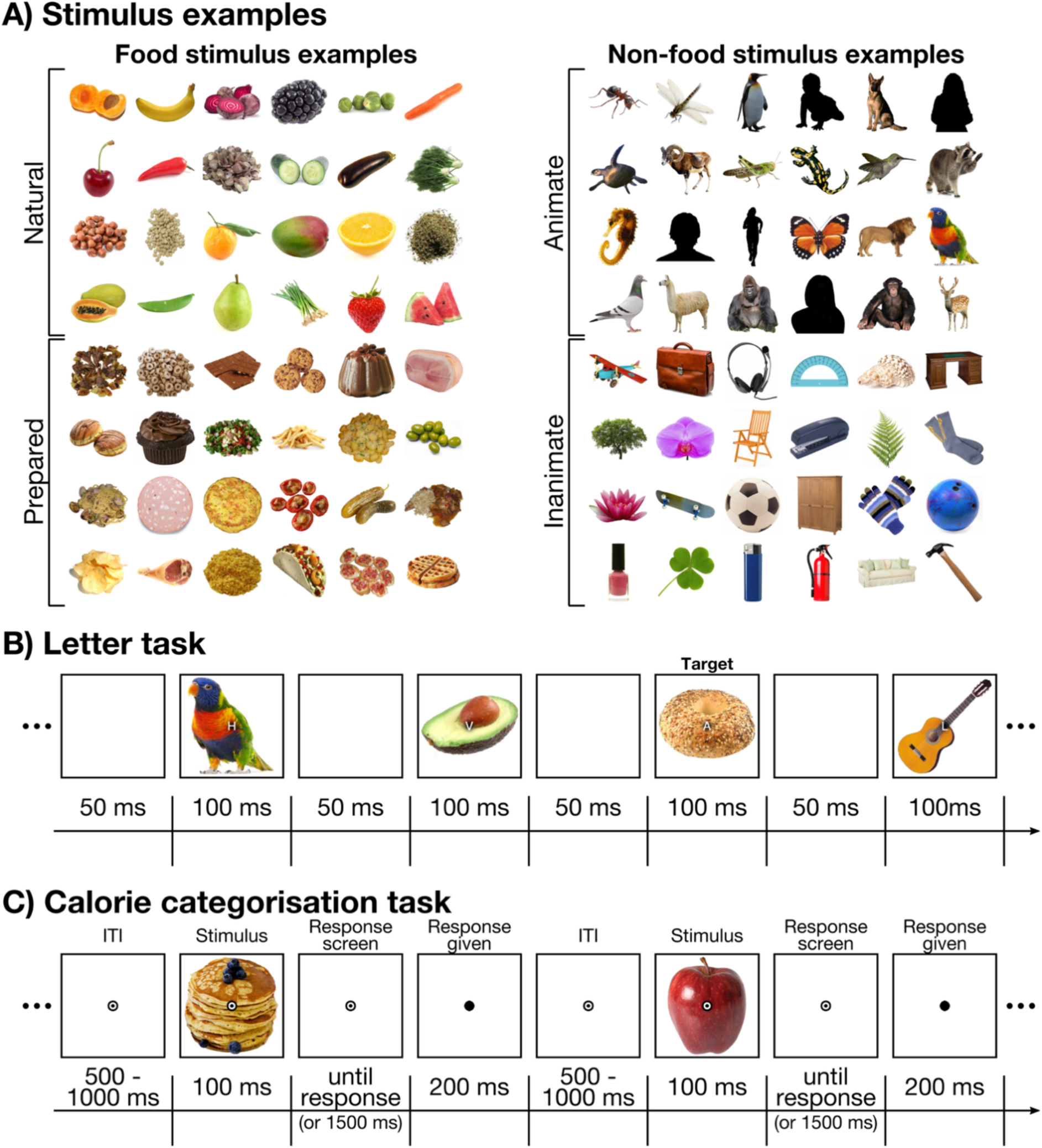
Example stimuli and EEG task overview. **A)** Examples of the stimuli that were used in the experiment. There were 140 food stimuli (examples displayed on the left) and 140 non-food objects (examples displayed on the right). Note the stimuli depicting humans have been replaced with silhouettes in this figure to comply with bioRxiv guidelines. **B)** An overview of the letter task. Participants monitored a stream of letters on the screen and pressed a button when they saw a vowel. The images behind the letters are not task relevant. The letters and images were on the screen for 100 ms, with a 50 ms blank screen between presentations. **C)** An overview of the calorie categorisation task. Participants were presented with a food image for 100 ms and were asked to rate the calorie content as higher or lower compared to bread. There was a 1500 ms response time-out.

#### 2.2.3. Tasks

The stimuli were presented in the centre of the computer screen on a white background, using the Psychopy library (Peirce et al., 2019) in Python. The participants were seated approximately 60 cm from the computer screen, and the stimuli were approximately 6.25 degrees of visual angle in size (256 x 256 pixels). The fixation bullseye was displayed at the centre and was approximately 0.61 degrees of visual angle (24 x 24 pixels) in size. Participants were instructed to keep fixation on the central bullseye during the task. The EEG experiment consisted of a letter task and a calorie categorisation task (see Figure 1B). There were 6 blocks in total, and each block consisted of the letter task followed by the calorie categorisation task. The experiment took approximately 40 minutes to complete.

In the letter task, participants saw a rapid stream of food and non-food images, with a simultaneously presented stream of letters, overlaid on top of the object images. Participants were instructed to ignore the objects and press a button whenever they saw a vowel (A, E, I, O, or U) in the stream of letters, as quickly and as accurately as possible. This means that the object images were not task relevant and attention was directed away from the stimuli and towards the letters.

Each of the 140 food images and 140 non-food images were presented three times per block and thus 18 times in total across the 6 blocks. Therefore, participants saw 5040 images in total. The order of images was randomised such that food and non-food images were intermixed and all items were presented before they were repeated a second and third time in each block. The object images with overlaid letters were presented at a rate of 6.67 Hz, where each stimulus was on the screen for 100ms, followed by a blank screen for 50 ms.

Each sequence contained 2-4 targets (vowels). Letters were presented in a randomised order, however, there were always at least 15 random letter presentations between two targets and the first 10 and last 10 letters within each sequence were never targets. Letters ‘Q’ and ‘Y’ were not used, as ‘Q’ looked too similar to ‘O’ at a rapid presentation rate and ‘Y’ could be confused for a vowel. Task performance was indicated by mean accuracy and reaction time. Mean accuracy was calculated per session, as the number of correct responses made within 1.5 seconds post target presentation, divided by the total number of targets seen. Mean reaction time in seconds was calculated per session, as the mean time of correct responses.

In the calorie categorisation task, participants saw images of food items and were asked to classify the foods as higher or lower in caloric density compared to bread as quickly and accurately as possible (Figure 1C). Specifically, they were asked to compare 100g of the presented food to 100g of bread in terms of the caloric density. The task was designed to ensure the stimuli were attended to without focusing on the specific dimensions of interest (flavour, appeal, arousal). We chose bread as a reference point as it is familiar to Australian participants and was close to the median perceived calorie rating in FRIDa, effectively dividing the set into ‘higher’ or ‘lower’ calorie categories. Participants’ responses to this task were not used in analyses. Participants completed this classification by pressing the left or right outer buttons of a button box. The buttons for “higher calories” and “lower calories” changed between blocks to minimise the effect of motor preparation and execution signals that could contaminate the EEG signal.

The 140 food stimuli were shown once per block in a random order, meaning each stimulus was shown 6 times in total across the 6 blocks of the classification task. Participants took voluntary breaks after every 20 trials to minimise fatigue. During these breaks, the instructions reappeared on the screen to remind the subject which button mapped to high or low calories. In each trial, the fixation point was shown on the screen for 500-1000ms to prevent participants from anticipating the stimulus onset. The food was shown for 100ms, then replaced by the fixation point again for 1500ms or until the participant responded. If the participant responded within 1500ms, the circle would change to black to indicate that their response was registered and the next trial would start. If no response was registered, a message saying “Too late!” appeared on the screen to encourage the participant to respond faster on the next trial.

### 2.3. Behavioural analysis

To determine whether our hunger manipulation was successful, we tested for a difference between sessions in the reported hunger level, time since the last reported meal, and time since the last reported food intake (including snacks). As a control, we tested for a difference between sessions in the reported thirst level and fatigue.

Participants also rated the appeal of each food item at the start of both EEG sessions, and we tested for a difference in reported food appeal across sessions. Previous work showed faster response times for high-fat (i.e. high-calorie) foods over low-fat foods in a visual search task when participants were hungry compared to when they were sated (Sawada et al., 2019). In addition, there is evidence participants choose high calorie over low calorie foods when hungry (Read & van Leeuwen, 1998; Tuorila et al., 2001). This suggests that hunger effects on appeal ratings could be stronger for high-calorie food items. We therefore split the appeal ratings based on the perceived caloric content, comparing the appeal of high calorie (top 25%) and low calorie (bottom 25%) foods across the fasted and fed conditions.

### 2.4. EEG Recording and Pre-processing

We continuously recorded the 64-channel EEG data at a sampling rate of 1000Hz using a BrainVision (GmbH, Herrsching, Germany) ActiChamp system. The electrode locations corresponded to the international 10-10 system of electrode placement (Oostenveld & Praamstra, 2001) with the reference at FCz. EEG data pre-processing was conducted offline using the EEGLAB toolbox (Delorme & Makeig, 2004) in MATLAB. We used minimal EEG pre-processing, following earlier work (Moerel, Psihoyos, et al., 2024). We used a high-pass and low-pass filter to remove signals below 0.1Hz and above 100Hz, respectively, and down-sampled the data from 1000Hz to 250Hz. We then created epochs for each trial, starting 100 ms before stimulus onset and ending 1200 ms after stimulus onset.

### 2.5. Decoding analysis

We used decoding to gain insight into the time-course of the following features: 1) edibility (food versus non-food object) and 2) food identity (e.g. hamburger versus pizza). We used CoSMoMVPA toolbox for Matlab (Oosterhof et al., 2016) for all decoding analyses, using linear discriminant analysis classifiers on the pattern of activation across all electrodes to distinguish between the classes of interest. We did this separately for each time-point and participant. Critically, we also did this separately for the sessions where the participant had fasted or eaten and then compared decoding accuracies across the two different sessions.

For the decoding of edibility information, we only used the data from the letter task for this analysis, as the calorie categorisation task did not have non-food object items. We used an exemplar-by-block cross validation approach, following previous work (Carlson et al., 2013; Grootswagers et al., 2019; Moerel, Psihoyos, et al., 2024), leaving out one pair of images (one randomly matched food and non-food object item), from one block as the test data. We repeated this analysis 10 times, with different random image pairings, and averaged the decoding accuracies across the repeats.

To determine *if* and *when* there was information about the identity of the food, we decoded pairs of food stimuli. For this analysis, we used the data from both the letter task, including the only the food items, as well as the data from the calorie categorisation task. We repeated the decoding analysis for each possible combination of two food stimuli and then averaged the decoding accuracy across the possible pairs. For the letter task we used a leave-one-block out cross-validation approach for each pair of stimuli, where all stimulus presentations from a single test block were left out in an iterative fashion. For the calorie categorisation task, we swapped the response button mapping between blocks to avoid motor response or preparation contamination of the EEG signal. We therefore tested the classifier on data from two blocks, one with each response mapping. We used all combinations of two blocks with opposite response mapping as the test blocks and averaged across the decoding accuracies.

### 2.6. Representational Similarity Analysis

We used Representational Similarity Analysis (RSA) (Kriegeskorte et al., 2008; Kriegeskorte & Kievit, 2013) to examine whether information about the flavour of food and current valence and arousal were represented in the brain. We calculated Representational Dissimilarity Matrices (RDMs) for the EEG data, representing the dissimilarity in the pattern of activation across the sensors evoked by two different food stimuli. We used the classification accuracy from the pair-wise decoding analysis described above as a measure of dissimilarity. We did this for all possible combinations of two food items, resulting in a 140 by 140 RDM (Figure 2A). We repeated this analysis for each time-point in the epoch, separately for each participant and session. We obtained three model RDMs, coding for 1) the flavour profile of the food, 2) the current appeal of the food, and 3) the current arousal of the food. Models for the current appeal and arousal were based on behavioural ratings obtained from the same EEG session. We then calculated partial correlations between the models and the EEG RDMs, for each time-point, participant, and session. To account for possible low-level visual differences between the stimuli, we partialled out three artificial neural network control models (V1, V2, and V4 layers of CORnet-S (Kubilius et al., 2019)), resulting in partial correlations between the EEG and the models.

**Figure 2.**
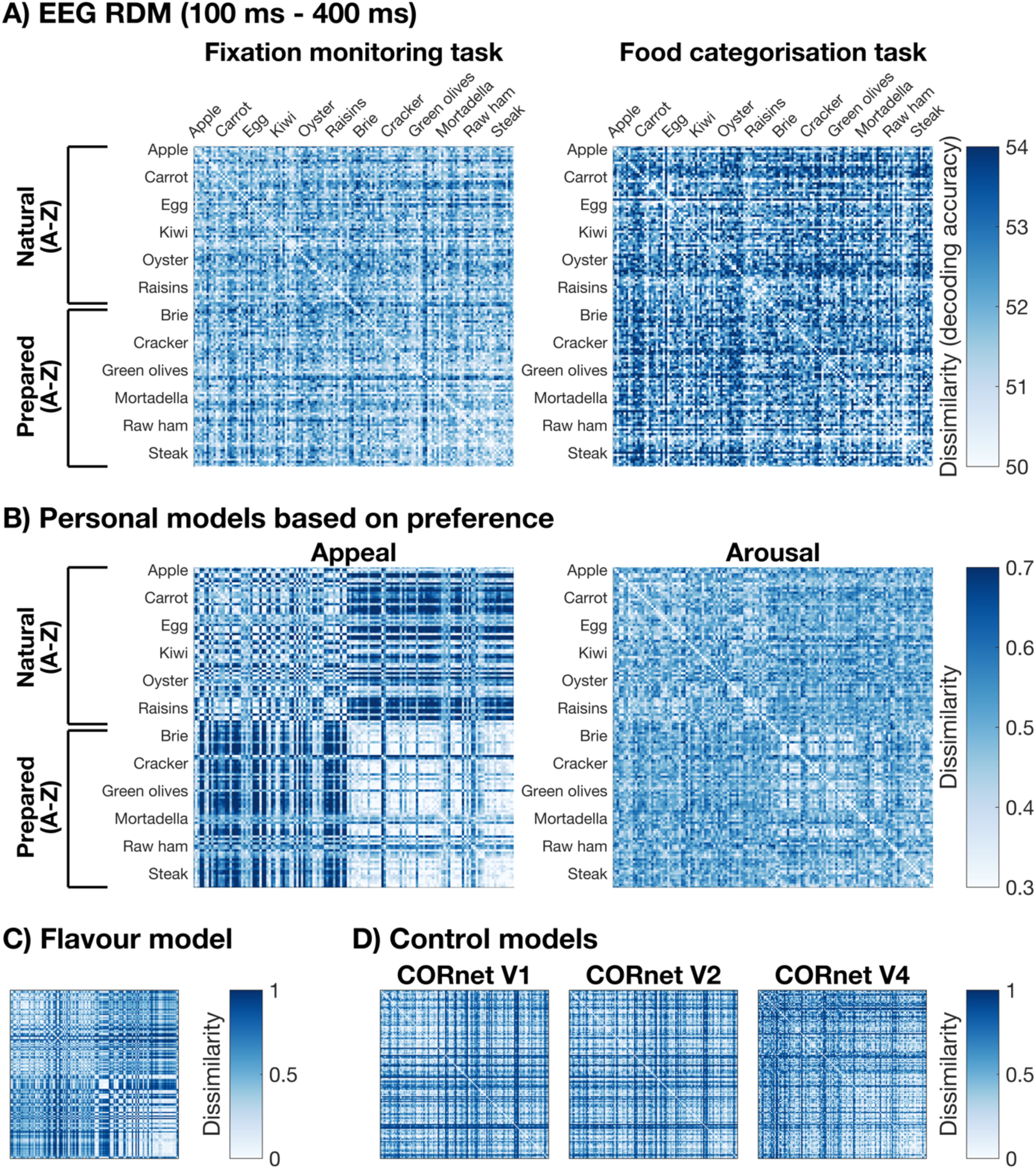
Representational Similarity Analysis overview. We made 140 x 140 Representational Dissimilarity Matrices (RDMs) representing the dissimilarity between all pairs of food images. **A)** The average RDM for the EEG data from the letter task (left) and calorie categorisation task (right). The EEG RDMs show the dissimilarity in the pattern of activation across the sensors, for each pair of stimuli. We make a unique RDM for each participant, session, and time-point. The RDMs plotted here show the mean across all participants and sessions, for the time window between 100 ms and 400 ms relative to stimulus onset. **B)** The individual appeal (left) and arousal (right) models. We made a personal model for each participant and session, based on behavioural ratings obtained at the start of the experiment. Mean RDMs across all participants and sessions are shown here. **C)** The general flavour model. This model was based on behavioural ratings from a different group of 84 participants. **D)** The three CORnet-S control models. These RDMs model the low-level visual differences between the stimuli.

#### 2.6.1. Flavour profile model

The model of the flavour profile (Figure 2C) was based on a different sample of 84 participants (mean age = 20.48, SD age = 2.60, age range = 18 – 33, 74 right-handed / 10 left-handed, 57 female / 26 male / 1 non-binary). Participants rated the 140 food stimuli on four different flavour dimensions: sweet, salty, sour, and bitter. Participants used a sliding scale to rate each of the food items on a single taste dimension per block, covering all taste dimensions across four different blocks. We obtained the flavour profile RDM by calculating the Euclidean distance between two pairs of food stimuli, on the pattern of responses across the four flavour dimensions. We did this separately for each participant and then averaged across participants to obtain a single flavour profile RDM.

#### 2.6.2. Personal appeal and arousal models

We calculated an individual appeal and arousal RDM for each participant and session (Figure 2B). Unlike the flavour profile model, the appeal and arousal models were based on behavioural data from the same participants that participated in the EEG session. Both models were based on the appeal rating, provided on a sliding scale from 0 to 100 at the start of each EEG session. The appeal RDM was obtained by calculating the absolute difference (dissimilarity) in the appeal rating between each combination of two food stimuli. The arousal RDM coded for the similarity in the emotional response that was elicited by the food, regardless of whether the food was considered appealing or unappealing. To convert the appeal rating into an arousal rating, we recoded the ratings as the absolute distance to the middle possible rating (50). We then calculated the arousal RDM in the same way as the appeal RDM.

#### 2.6.3. Visual control models

To control for possible low-level differences between stimuli, we used three artificial neural network control models (Figure 2D). We used the V1, V2, and V4 layers of CORnet-S (Kubilius et al., 2019), designed to reflect the response of visual areas V1, V2, and V4 respectively. We obtained the neural network responses to all stimuli and created the RDM by calculating the cosine distance in response between each pair of images. We then calculated the partial correlations between the EEG and model RDMs by partialling out the three control RDMs.

### 2.7. Exploratory analyses

#### 2.7.1. Generalisation of appeal and arousal over sessions and participants

We explored whether the information about personal appeal and arousal generalises 1) across time and hunger state, and 2) across different participants. To test whether information about appeal and/or arousal generalises across hunger states and time, we calculated the correlations between the EEG data, and the appeal and hunger models based on the behavioural data from the other session. If this information generalises across sessions, that suggests we are tapping into preferences that are stable across time. If not, this suggests that the appeal/arousal information is driven by the temporary state of the participant. We also tested whether information about appeal and/or arousal generalises across participants. If so, this would suggest the information is driven by general preferences across the population, rather than personal preferences. For this analysis, we calculated the correlations between the EEG data, and the appeal and hunger models based on the behavioural data from a randomly matched participant in the same hunger state condition. This means the hunger level was matched between participants. We repeated this analysis 10 times, with different random matches between participants, and then averaged the correlation across the 10 repeats.

#### 2.7.2. The effect of appeal on the coding of edibility information

We also explored whether there was an effect of appeal on the coding of edibility information in the brain. We used the results from the edibility decoding analysis described above, training and testing on all 140 food stimuli and 140 non-food object items. As described above, this analysis was only performed on EEG data from the letter task. For this exploratory analysis, we split the decoding accuracies based on whether the presented item was high appeal (top 25%) or low appeal (bottom 25%). We collapsed across hunger state conditions. We then compared the edibility decoding for high appeal and low appeal food items.

### 2.8. Statistics

We used Bayesian statistics (Dienes, 2011; Kass & Raftery, 1995; Morey et al., 2016; Rouder et al., 2009; Wagenmakers, 2007), as they offer an intuitively interpretable measure of evidence for or against hypotheses, allow us to determine when there is sufficient evidence to draw conclusions in either direction, and enable us to accept the null hypothesis. The statistics were implemented through the Bayes Factor R package (Morey & Rouder, 2018), to determine whether there was evidence for above chance decoding or partial correlations above 0. We applied Bayesian t-tests at each time-point. All statistical analyses were applied at the group level, and we used the same analyses for the letter task and calorie categorisation task. For the decoding analysis, we tested whether there was evidence for the chance decoding (null hypothesis) or the above chance decoding (alternative hypothesis). For the RSA analysis, we tested whether there was evidence for a partial correlation of 0 (null hypothesis) or above 0 (alternative hypothesis). We used a point null for the null hypothesis and a half-Cauchy, directional, prior for the alternative. The prior was centred around chance (d = 0, chance decoding or a partial correlation of 0), and we used the default prior width of d = 0.707 (Jeffreys, 1998; Rouder et al., 2009; Wetzels et al., 2011). We excluded small effect sizes, that may occur by chance under the null hypothesis, by excluding the interval between d = 0 and d = 0.5 from the prior (Teichmann et al., 2022).

We also applied Bayesian t-tests for our behavioural ratings. We followed the same steps as described above with the following differences. To obtain Bayes Factors for the differences between the fasted and fed conditions, we subtracted the ratings for the fasted and fed conditions and performed Bayesian t-tests against 0. We used a full-Cauchy rather than a half-Cauchy prior, to make the test 2-tailed. We excluded the interval between d = −0.5 and d = 0.5 from the prior for consistency with the t-tests described above.

## 3. RESULTS

### 3.1. Behavioural results

At the start of both EEG sessions, participants answered questions to determine their level of hunger, thirst, and fatigue, as well as the time since their last food intake and full meal. The results suggest that the hunger manipulation was successful, resulting in higher reported hunger levels (BF = 6.75 × 10^5^), a longer time since the last full meal (BF = 9.35 × 10^8^), and a longer time since the last food intake including snacks (BF = 1.30 × 10^14^) when participants were asked to fast (Figure 3A). Control analyses showed evidence for no difference between sessions in the reported thirst level (BF = 0.05) or fatigue (BF = 0.01).

**Figure 3.**
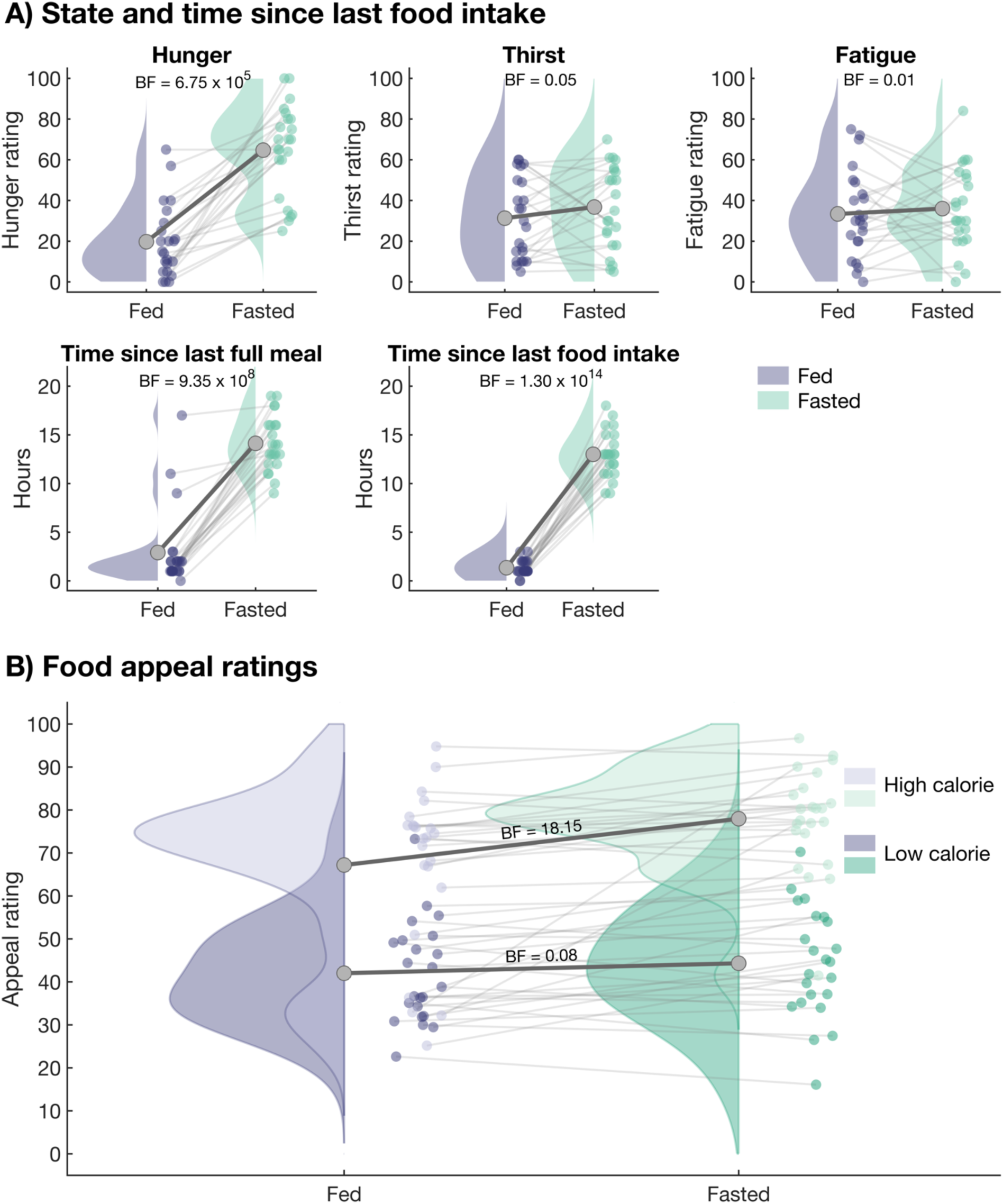
Behavioural results. **A)** The responses to the questionnaire taken at the start of each EEG session, in the session where participants had eaten (purple) or were asked to fast (green). Dots show individual participants, and the thick grey line shows the mean across participants. The top panel shows the rating for the hunger level (left), thirst (middle), and fatigue (right). The bottom panel shows the reported time since the last full meal was consumed (left) and the time since any food, including snacks, was consumed (right). **B)** Participants also rated the appeal of each food items at the start of each EEG session, with food appeal ratings for the session where participants had eaten (purple) or were asked to fast (green) shown here. We split the data based on the perceived calorie content (Foroni et al., 2013), showing high calorie foods (top 25%) in light colours and low calorie foods (bottom 25%) in dark colours. Plotting conventions are the same as Figure 3A.

We compared the food appeal ratings of high calorie (top 25%) and low calorie (bottom 25%) foods across the fasted and fed conditions (Figure 3B). For the low-calorie foods, we found no evidence for an effect of hunger state (BF = 0.08). However, there was an effect of hunger state for the high calorie foods (BF = 18.15), with an increased appeal of high calorie foods when the participants were hungry compared to sated.

### 3.2. No effect of hunger on the neural encoding of food-related features

We investigated whether the hunger state of the participant influences the processing of food stimuli in the brain. We hypothesised that if previous univariate results reflect a general hunger-driven enhancement of the processing of food-related information, we should find stronger multivariate encoding of these features when participants had fasted. Specifically, we investigated information about edibility, food identity, flavour, personal appeal, and arousal across fasted and fed conditions.

First, we used the EEG data from the letter task to decode whether a stimulus item was a food or non-food object. This allowed us to investigate the time-course of edibility information when participants were asked to fast or had just eaten. The results are plotted in Figure 4. There was evidence for edibility information from 84 ms after stimulus onset for both the fasted condition and the fed condition. However, there was evidence for no difference between the fasted and fed condition for most of the time-course, and no time-points showed strong evidence for a difference.

**Figure 4.**
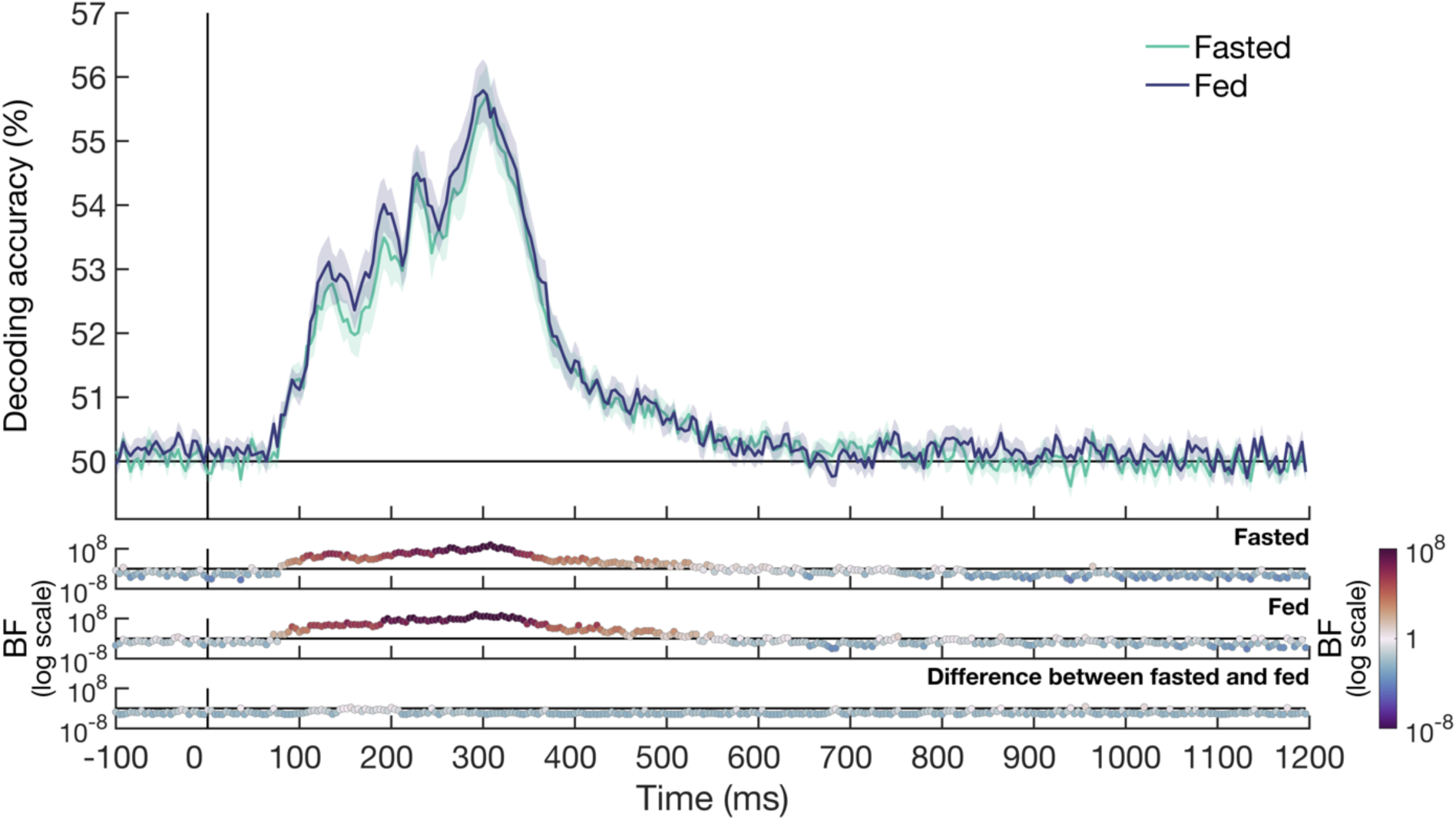
The time-course of edibility decoding for the fasted and fed conditions. We used a linear classifier on the EEG data from the letter task to distinguish between food items and non-food objects. The decoding accuracy over time is shown for the fasted condition (green) and the fed condition (purple). Shaded areas around the plot line show the standard error of the mean. Theoretical chance is at 50% decoding accuracy. The Bayes factors (BFs) are shown on a logarithmic scale below the plot for the fasted condition (top), fed condition (middle) and the difference between the two conditions (bottom). BFs above 1, showing evidence for above chance decoding, are plotted in warm colours and BFs below 1, showing evidence for chance decoding, are plotted in cool colours.

We also investigated whether the hunger state of the participant could influence the coding of more fine-grained information about the food. To investigate this question, we decoded the identity of the food in a pair-wise way. Figure 5 shows the food identity decoding for the calorie categorisation task (Figure 5A) and the letter task (Figure 5B). For the calorie categorisation task, we found evidence for food identity decoding from 96 ms after stimulus onset in the fasted condition, and 100 ms after stimulus onset in the fed condition. There was evidence for no difference in food identity decoding between the fasted and fed conditions for most time-points, with a few time-points showing insufficient evidence for either hypothesis. The pattern of results for the letter task was similar, with evidence for food identity decoding from 88 ms onwards in the fasted condition, and from 96 ms after stimulus onset in the fed condition. Again, there was evidence for no effect of hunger on food identity decoding for most time-points, with insufficient evidence found for a few time-points.

**Figure 5.**
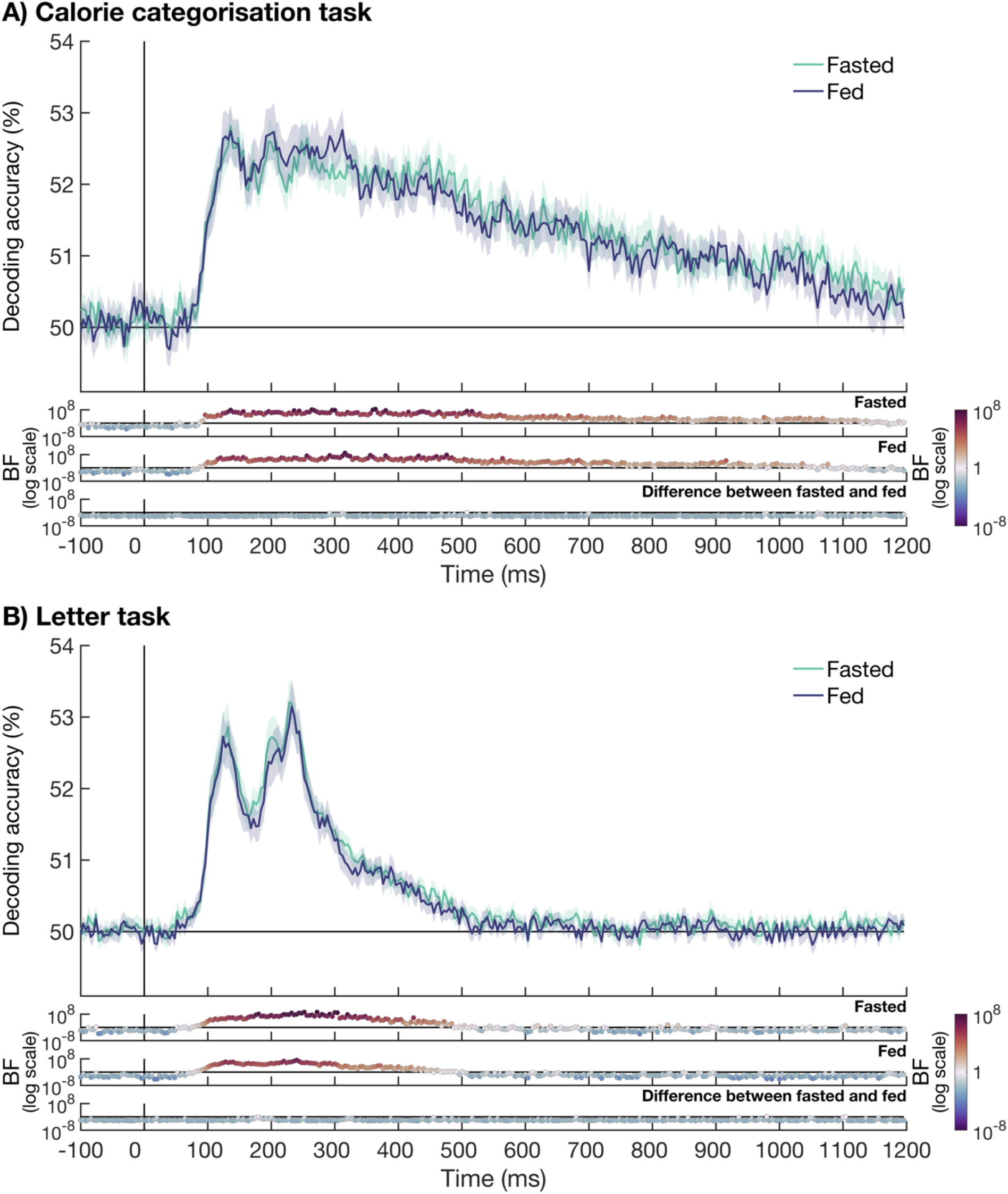
The time-course of food identity decoding for the fasted and fed conditions for the calorie categorisation task (A) and letter task (B). We used a linear classifier on the EEG data to distinguish between two food items in a pair-wise way. Plotting conventions are the same for both plots. The decoding accuracy is plotted over time for the fasted condition (green) and fed condition (purple). Theoretical chance is at 50% decoding accuracy, and shaded areas around the plot line show the standard error of the mean. The Bayes factors (BFs) are shown below the plot on a logarithmic scale. The top plot shows the BFs for the fasted condition, the middle plot for the fed condition, and the bottom plot for the difference between the two conditions. BFs above 1 are plotted in warm colours and BFs below 1 are plotted in cool colours.

We used RSA to gain further insight into the effect of hunger on the coding of information about the flavour profile, appeal, and arousal of food. The flavour profile was based on data from a different group of participants, i.e., one general flavour profile model. The appeal and arousal models were based on data from the EEG participants, obtained in the same session, with a unique appeal and arousal model made for each participant and for each session. We calculated partial correlations between the EEG data from the calorie categorisation task or the letter task and the models, controlling for three low-level visual CORnet models. The data, separated for the fasted and fed condition, can be found in Supplementary Figure 1. Consistent with the findings from the decoding analysis, there was no evidence for an effect of hunger on information about the flavour profile, appeal, and arousal of food, for either the calorie categorisation task or the letter task. Most time-points in all tests showed evidence for no difference between the partial correlation from the fasted and fed conditions. Together, these results suggest that even though the brain processes these different food-related features, whether an object is a food, the identity of the food, flavour, personal appeal, and arousal, this information is not affected by the hunger state of the participant.

### 3.3. Attention-independent processing of some, but not all, food-related features

This study aimed to assess whether focused attention was necessary for the brain to encode information about food identity, flavour, personal appeal, and arousal. We hypothesised that if certain food-related features are represented in the brain regardless of focused attention or task relevance, then these representations occur as a consequence of simply perceiving the food stimulus. Such attention-independent representations would suggest that these features may be processed automatically. Our results showed the neural signal contained information about identity of the food stimulus when the stimuli were task-relevant (Figure 5A) as well as when attention was focused on an orthogonal task (Figure 5B). Figure 6 shows the time-course of information about the flavour profile, appeal, and arousal of food. For the calorie categorisation task, there was evidence for information about the flavour profile from approximately 192 ms onwards. There was also information about the appeal and the arousal of the food item, but this information emerged later, around 520 ms and 464 ms after stimulus onset respectively. The letter task, where the food items are not task-relevant, shows a different pattern of results. For this task, we find information about the flavour profile from approximately 212 ms onwards. However, there is no consistent information about either appeal or arousal, with evidence for no partial correlation for most time-points. Together, these results show that there is information about the food identity and flavour profile of the food, regardless of the task. However, there was only information about personal appeal and arousal in the calorie categorisation task, when the task is task relevant, and the stimuli are presented at a slower rate, suggesting that focused attention is needed for these features to be encoded.

**Figure 6.**
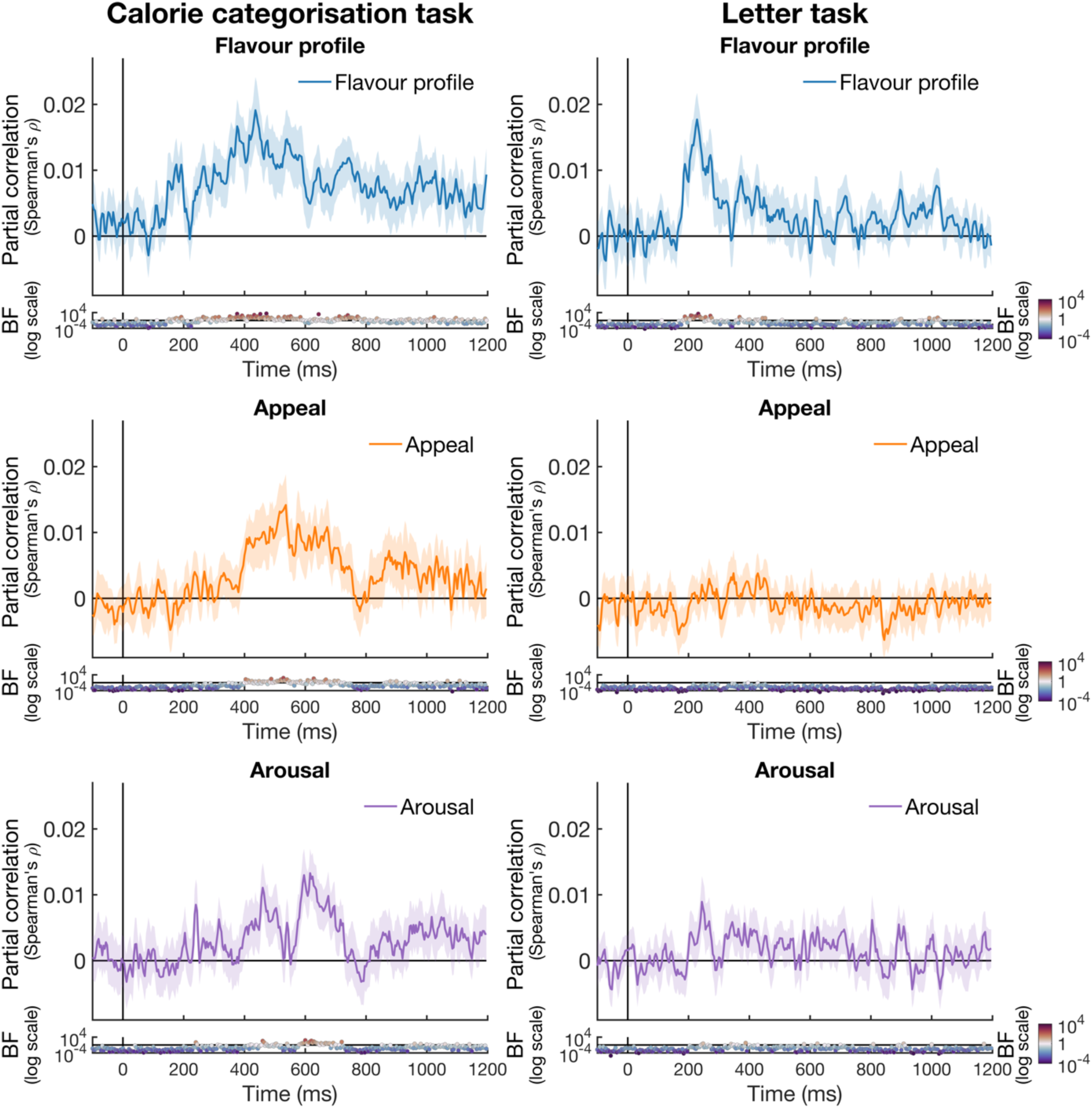
The time-course of partial correlations between the EEG data and the flavour profile, appeal, and arousal models. We calculated partial correlations between the EEG data for the calorie categorisation task (left) and the letter task (right) and the models. The top panels show the partial correlation with the flavour profile model, the middle panels show the partial correlation with the individual appeal models, and the bottom panels with the individual arousal models. To account for possible low-level visual differences between the stimuli, we partialled out three CORnet control models from the correlations. The shaded areas around the plot lines show the standard error of the mean. The Bayes Factors (BFs) are shown below the plot. BFs above 1 are plotted in warm colours and BFs below 1 are plotted in cool colours.

### 3.4. The time-course of appeal and arousal information in the brain

The final aim of this study was to assess the time-course of neural information about appeal and arousal, obtained through individual ratings in the same session. We hypothesised that personal appeal and arousal would be encoded in the neural signal, and that is reflects highly personal, rather than more general, preferences. Therefore, in addition to determining whether there was neural information about appeal and arousal based on the behavioural ratings from the same session, we included a control analysis where we focused on data from the calorie categorisation task to further explore whether this information about appeal and arousal generalised to 1) appeal/arousal ratings from the other session for the same participant and 2) appeal/arousal ratings from other participants. If there is a correlation between the EEG and the personal appeal/arousal model within participants across sessions, this suggest that this information reflects long term food preferences of that individual, that are stable over time. If there is a correlation between the EEG and the personal appeal/arousal model across participants, this suggest that this information reflects universal food preferences that generalise across the population.

Figure 7 shows the generalisation of appeal and arousal information. As described in the previous section, there was information about the appeal of the food item from approximately 520 ms after stimulus onset when we used the individual appeal model that was matched for the participant and session. When using the appeal model for the same participant, but the other session, we found no reliable evidence for information about appeal. Although visual inspection shows there was a trend for a correlation, showing a similar time-course as the EEG correlation with the model based on the same session, the BFs show the evidence for this correlation is anecdotal. We find a similar pattern of results for the correlation with a model from a randomly matched participant. Visual inspection shows a small correlation with a similar time-course as the matched appeal model, but this is not supported by the BF evidence. For the arousal model, we previously found a correlation between the EEG RDM and the individual arousal model that was matched for the participant and session from approximately 464 ms after stimulus onset. There was no evidence for a correlation between the EEG and the arousal model from the same participant but other session, with most time-points showing evidence for the null. In addition, the model based on a random matched participant from the same session also did not correlate with the EEG, most time-points showing evidence for the null. However, there were 2 consecutive time-points with BF > 10 around 150 ms after stimulus onset. Neighbouring time-points did not show evidence for a correlation above 0, suggesting this is likely driven by noise. Together, these results show that only the matched appeal and arousal within the same session models reliably correlate with the EEG RDMs. However, there is a possibility that appeal could generalise to the other session and perhaps even across participants, but this evidence is merely anecdotal.

**Figure 7.**
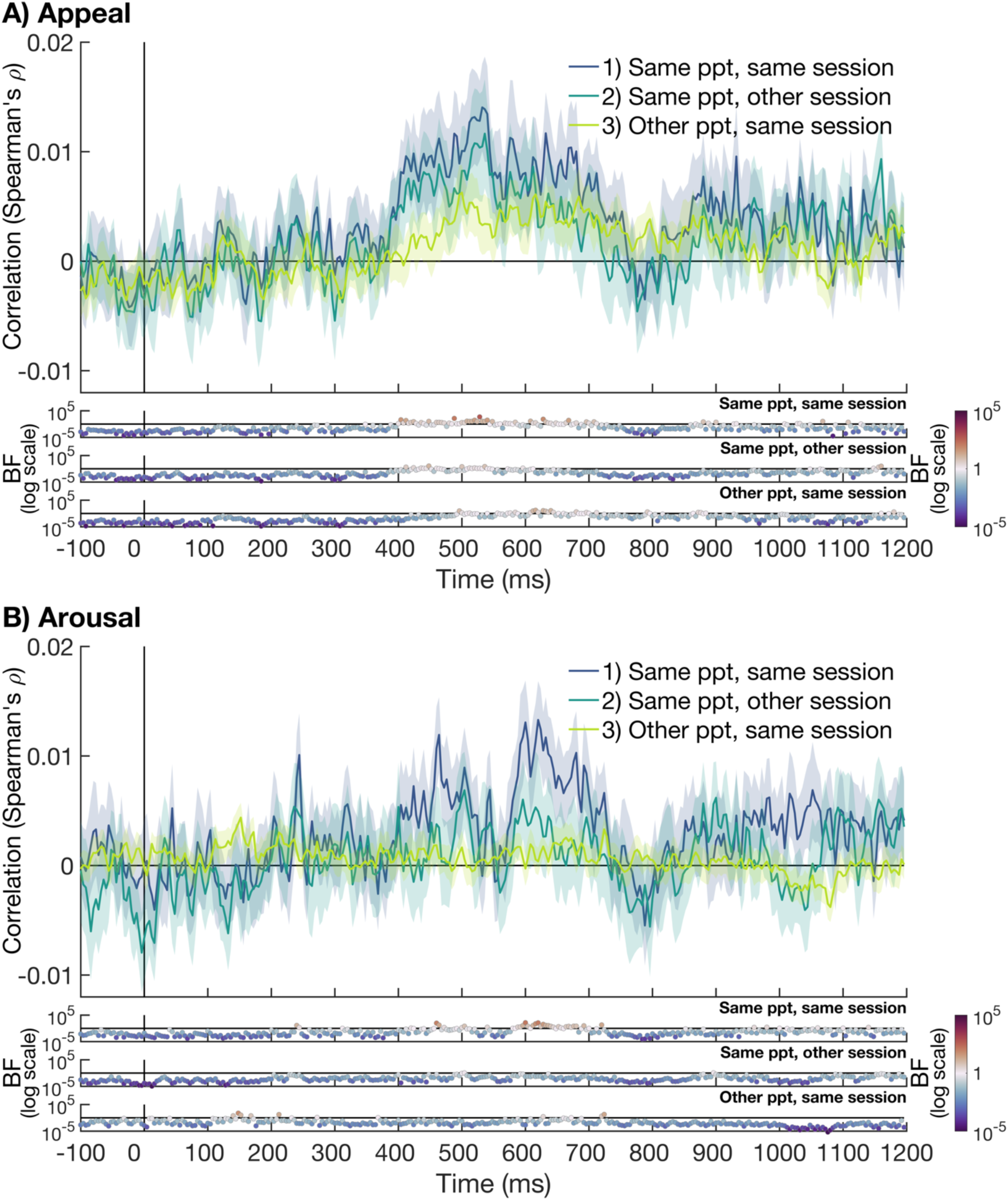
the generalisation of food appeal (A) and arousal (B) information across sessions and participants. We explored the correlation between the EEG for the calorie categorisation task and the appeal and arousal models, that were based on 1) the same session and participant (dark blue), the same participant but the other session (teal), a randomly matched participant but the same session (light green). The shaded areas show the standard error of the mean. The Bayes Factors (BF) are shown below the plot on a logarithmic scale. BFs above 1 are shown in warm colours and those below 1 are shown in cool colours.

### 3.5. Exploratory analysis: The neural representation of edibility is mediated by personal appeal

We explored if the representation of edibility information, whether an item is a food or not, is affected by the appeal or arousal of the food. For this analysis, we used the same edibility decoding data presented in section 3.2, collapsed the edibility decoding accuracy across the fasted and fed conditions, and split the results based on whether the food item had high appeal (top 25%) or low appeal (bottom 25%) or high or low arousal (top versus bottom 25%). We did not include the results for the middle 50% of food items and the non-food items. The EEG data were obtained from the letter task, where the objects were not task relevant, and the food items were therefore likely not within the focus of attention. The results are shown in Figure 8. When splitting the results based on appeal, there was information about whether an item was edible from 92 ms onwards for the high appeal items and 84 ms onwards for the low appeal items. However, after approximately 176 ms, there was an effect of appeal, with stronger edibility decoding for the high appeal compared to low appeal items. When splitting the data by arousal, we found edibility decoding from 88 ms onwards for the high arousal items and after 92 ms for the low arousal items. We did not find an effect of arousal on edibility decoding, with most time-points showing not enough evidence either way, or evidence for no difference between high and low arousal. Together, these results suggest that edibility information in the brain is affected by the appeal, but not arousal, of the food stimulus.

**Figure 8.**
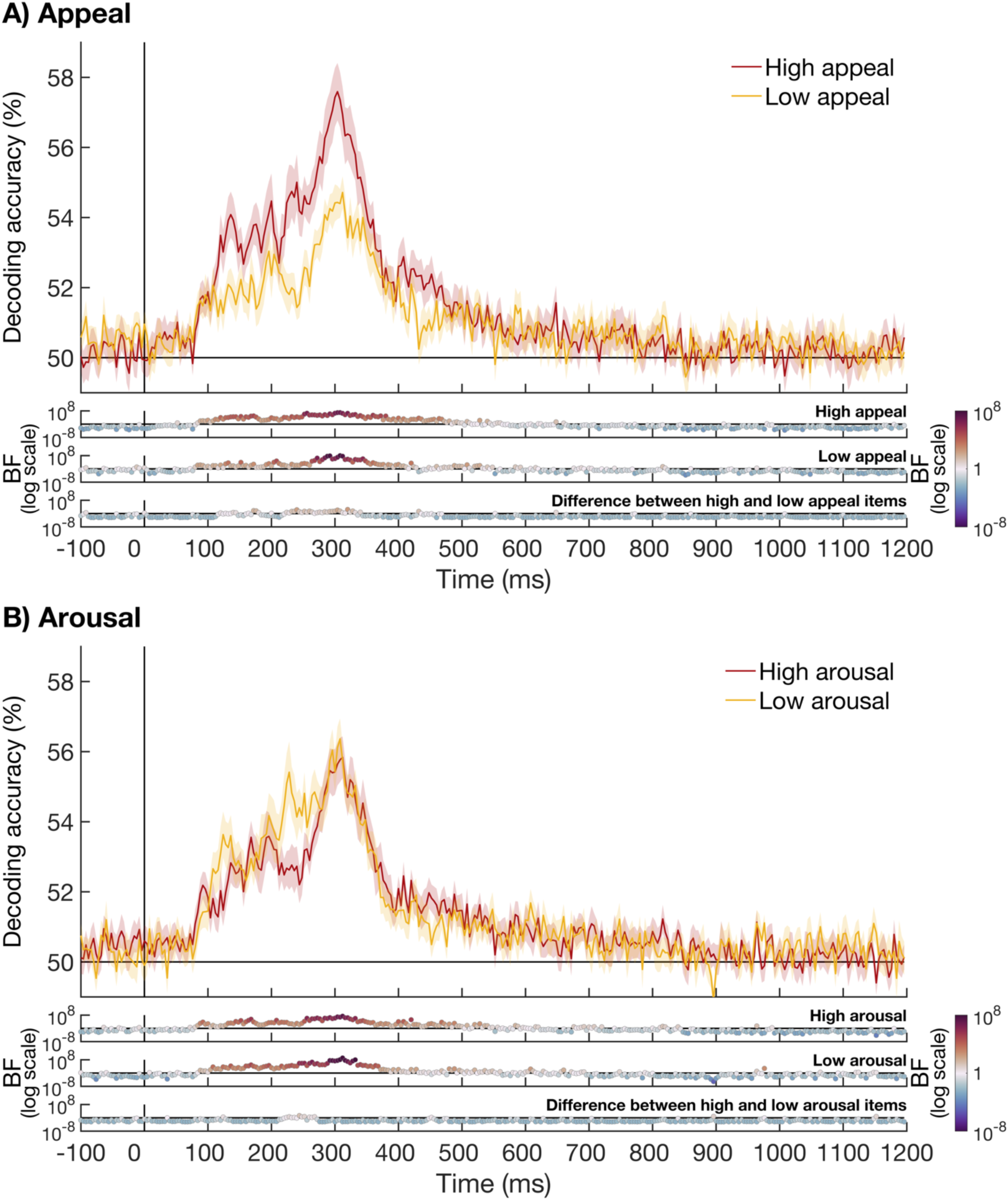
The time-course of edibility decoding, split by appeal (A) or arousal (B) level of the food items. **A)** We split the edibility decoding results, presented in Figure 4, based on whether the food item had high appeal (top 25%) or low appeal (bottom 25%). We excluded the results for the middle 50% of food items and non-food items. The decoding data for high appeal items is presented in red, and the data for the low appeal items is presented in yellow. Shaded areas around the plot lines show the standard error of the mean. The Bayes Factors (BFs) are presented below the plot, with BFs above 1 shown in warm colours and BFs below 1 in cool colours. **B)** shows the decoding data split by high arousal (top 25%) in red and low arousal (bottom 25%) in yellow. All plotting conventions are the same as in A.

## 4. DISCUSSION

In this study, we investigated 1) whether the hunger state of the participant influenced the neural representation of different food-related features, 2) whether focused attention, manipulated through task-relevance, was needed for the coding of food information, and 3) whether information about personal appeal was reflected in brain activity. We hypothesised that if hunger drives a general enhancement of food-related features, fasting should enhance the neural encoding of edibility, food identity, flavour, appeal, and arousal information in the brain. Additionally, we hypothesised that if food-related features are encoded independently of task demands, in line with automatic processing, their neural representations would persist without focused attention. Finally, we hypothesised that individual-specific ratings of personal appeal and arousal would be reflected in the neural signals, indicating encoding of highly personal food preferences.

Our EEG data showed that information about edibility (food versus non-food) and the identity of the food items (e.g., hamburger versus pizza) emerged early, before 100 ms. Critically, the hunger state of the participant did not affect the coding of this information, with no reliable difference between when the participant had fasted or eaten. Similarly, there was no effect of hunger on the coding of flavour, appeal, or arousal information. We found evidence that the representation of information about the flavour profile, emerging around 200 ms after stimulus onset, was present regardless of whether the food stimulus was task relevant or not. Information about the personal appeal and valence of the food emerged later and was only present for the calorie categorisation task, where the stimuli where task relevant and presented at a slow rate. In addition, the results showed that there was no reliable evidence that information about either appeal or arousal transferred to the other session or participants, although there was a trend. This suggests that appeal and arousal information is driven, at least in part, by the temporary state of the participant.

### 4.1. No effect of hunger on the neural processing of food-related visual information

Our results show no effect of hunger state on the representation of information about edibility, food identity, flavour profile, personal appeal, or personal valence, with strong evidence for no effect of hunger in most time-points. Previous work has shown evidence that hunger mediates neural responses to food items, using fMRI (Charbonnier et al., 2018; Cheng et al., 2007; Führer et al., 2008; Holsen et al., 2005; LaBar et al., 2001; Mohanty et al., 2008) and EEG (Ilse et al., 2020; Nijs et al., 2010; Stockburger et al., 2008, 2009; Sultson et al., 2019). There are several possible explanations for this apparent difference in findings between these previous studies and our work. First, previous studies used univariate methods to assess the effect of hunger on the neural activation and are therefore not able to discern whether the coding of information is affected by hunger. It is possible that the differences in activation, observed in previous work, are driven by other processes such as increased inhibitory control or differences in reward value when participants had fasted. For example, Charbonnier and colleagues (2018) used fMRI to show increased activation in the right dorsolateral prefrontal cortex when participants viewed high calorie images in a hungry state, which was interpreted as evidence for increased inhibitory control when participants had fasted. This could result in overall differences in activation without affecting the pattern of neural activation evoked by seeing the stimulus.

An alternative explanation for a lack of hunger effect is that our hunger manipulation was not successful, as the overnight fasting manipulation may have been too subtle. We relied on self-reported ratings of hunger rather than biochemical measures of hunger, and no physiological measures were used to confirm our manipulation was successful. However, it is important to note that all included participants reported complying with the instructions regarding food intake. In addition, there was evidence for a difference in reported hunger levels, hours since the last full meal, and hours since the last food intake (including snacks) between the fed and fasted conditions. There was also behavioural evidence showing an increase in the reported appeal for high-calorie food items when participants had fasted compared to when they had eaten. This difference was not observed for the low-calorie items, suggesting that our hunger manipulation was effective.

Another explanation for a lack of an effect of hunger is the stimulus set we used. There is some evidence that the stimuli that are chosen can influence whether hunger effects are observed. Using a visual mismatch negativity paradigm in EEG, Sultson and colleagues (2019) showed an effect of hunger between 100 ms and 220 ms after stimulus onset for high fat savoury stimuli. Importantly, there was no effect of hunger for the high fat sweet stimuli, highlighting the importance of stimulus choice. In line with this, other work has suggested hunger effects could be mediated by the motivational saliency of the stimuli. Piech and colleagues (2009) used fMRI to show enhanced responses to high compared to low appeal foods when the participant was hungry, but not when the participant had eaten. In addition, the calorie content of the food has been found to mediate hunger effects (Frank et al., 2010; Goldstone et al., 2009; Siep et al., 2009). Our study used a large stimulus set of 140 diverse food images, covering both high and low-calorie food of various appeal. It is therefore possible that the choice of stimuli can influence whether hunger effects are found.

Another possibility is that hunger effects were masked by our diverse sample of participants. For example, it is possible that collapsing results across men and women obscured hunger effects due to differences between groups. Previous fMRI work shows mixed findings, with some studies showing an interaction between gender and hunger (Frank et al., 2010), and other studies showing no interaction (Uher et al., 2006). However, a range of studies have found effects of hunger on the neural response to food stimuli when using samples of both women and men, in fMRI (Charbonnier et al., 2018; Cheng et al., 2007; Holsen et al., 2005; LaBar et al., 2001; Mohanty et al., 2008) and EEG (Ilse et al., 2020; Stockburger et al., 2009). This shows that evidence for hunger effects can be found when including both male and female participants. Another factor that could potentially mediate hunger effects is BMI (Nijs et al., 2010). Findings about the contribution of Body Mass Index (BMI) have been mixed (Ziauddeen et al., 2012). Some fMRI work showed that BMI can mediate neural responses to food (Volkow et al., 2011), whereas other fMRI work used multivariate analyses to show neural representation of visually presented food were not affected by BMI (Pimpini et al., 2022). Most of our participants had a BMI in the healthy range (18.5 – 24.9), two participants were underweight (BMI = 17.31 and 18.19), and three participants were overweight (BMI ranging between 26.35 and 27.98). No obese participants (BMI of 30 or higher) were included in the sample. As most participants had a BMI within the healthy range, it is unlikely this can explain the lack of an effect of hunger. Finally, most participants (20 of the 23) reported not being on a diet, with one participant reporting a pescatarian diet, one reporting a gluten free diet, one reporting an intermittent fasting diet. It is therefore also unlikely that differences in diet could have caused the lack of a hunger effect.

### 4.2. Dissociating attention-independent and attention-dependent food representations

Our results showed there was evidence for coding of both food identity (e.g., hamburger versus pizza) and flavour profile (e.g. savory versus sweet), regardless of whether the food items were task relevant. However, information about the current personal appeal and arousal was only present for the categorisation task, where the food stimuli were task relevant. Our findings show that focussed attention is needed for information about personal appeal and arousal to be encoded by the brain, whereas information about the food identity and flavour profile may be coded automatically. A key strength of our study is in testing which factors affect perceptual processing of food stimuli, under both focused and diverted attention, even without explicit value-based decision-making, simply upon seeing the stimuli. However, the stimuli in the calorie categorisation task were also presented at a slower rate compared to the letter task. It is therefore also possible that the faster speed of the letter task caused earlier masking (Robinson et al., 2019), limiting processing of the stimuli.

Flavour-related information emerged at approximately 200 ms after stimulus onset. It was encoded regardless of whether attention was directed towards the stimulus, although the response was less sustained over time when the food stimulus was not attended. The time-course of flavour-related information (∼200 ms relative to stimulus onset) is in line with top-down processes representing information about flavour, possibly reflecting semantic associations. Although we cannot fully rule out the role of visual heuristics, partialling out visual control models shows the flavour-related information is separate from visual stimulus features (e.g. colour implying sweetness).

### 4.3. Neural coding of food appeal reflects personal and dynamic preferences

Our results showed that there was information about personal appeal and arousal with regard to food stimuli during the categorisation task. There was no strong evidence for information about the appeal and arousal of food items generalising to a different session with the same participant, or across participants. The finding that the preference model from the same session explained variance in the EEG data, whereas there was only anecdotal evidence for the preference model from the other session, suggests that the information about appeal and valence was, at least partly, driven by the temporary state of the participant. Some preferences that are more stable over time might also have contributed to the information, driving the trend of shared appeal and arousal information across sessions. Although there was a trend suggesting some transfer between session and, to a lesser extent, participants, the current study does not have the power to statistically assess this. Overall, our study reconciles the different findings from previous work (Chae et al., 2025; Moerel, Psihoyos, et al., 2024; Schubert et al., 2021). Whereas several previous studies found that the brain carries information about the subjective pleasantness or ‘tastiness’ of food, using EEG (Chae et al., 2025; Schubert et al., 2021) and fMRI (Avery et al., 2025), our previous EEG work showed valence and arousal models based on a different group of participants did not explain the neural data (Moerel, Psihoyos, et al., 2024). Our current findings suggest that personal appeal and temporary cravings, rather than general preferences across the population, are a better predictor of the brain’s encoding of the food stimuli that did not include highly unappetising items. However, Chae and colleagues (2025) did find evidence that models of positive valence, negative valence and tastiness, based on one group of participants, could explain variance in the EEG data from another group of participants. A likely explanation, as suggested by Chae and colleagues (2025), is that their study included items that were specifically chosen to edible but highly unappetising. It is likely that agreement across the population about these unappetising items was high, thus enhancing between-subject model generalisation. In contrast, no highly unappetising items were included in our current and previous work, suggesting that any coding of appeal would be driven by more fine-grained, highly personal, and dynamic preferences in this case.

We further explored whether there was an effect of personal appeal on the coding of edibility information in the brain. Our results showed evidence for stronger edibility decoding in the brain for food items that were highly appealing compared to foods that were low in appeal. No such link was observed for arousal ratings. It is possible that the observed link between edibility decoding and subjective appeal is mediated by the personal experience with the food. It is reasonable to assume that the participants are more likely to consume the foods on a regular basis that are appealing to them (top 25%) compared to the foods that are unappealing (bottom 25%). This means participants might be more familiar with the appealing compared to unappealing items. In other words, the edibility decoding could be partly driven by “is it a food that I would eat” rather than “is it a food”. We did not ask participants about how often they consumed each of the food items in the stimulus set, which means it is not possible to further investigate the role of personal experience in edibility decoding.

### 4.4. Limitations and future directions

Our results suggest that hunger does not enhance the representation of visually presented food. Future work could further validate this finding by mapping effects across diverse stimulus sets and participant characteristics, and by using stronger physiological measures of hunger. Mapping effects across diverse stimuli and participants could further elucidate whether the absence of hunger effects on the *pattern of neural activation* in response to food images is a general effect, or whether this could be mediated by stimulus or participant characteristics. In addition, our results show that focused attention is not needed for brain to represent information about food identity and flavour, whereas this is the case for personal appeal and arousal. Although these results suggest that food identity and flavour may be processed automatically, future work is needed to further verify this. In addition, future work could further explore the processes that underlie the neural representation of flavour, evoked by seeing a stimulus (Avery et al., 2021). Finally, our exploratory results suggest that neural representations of appeal and arousal are highly personalised and do not fully generalise across sessions. In addition, appeal may mediate the representations of other food-related stimuli. Further work is still needed to disentangle the contribution of immediate appeal, potentially driven by temporary cravings, long-term food preferences, and general food preferences across the population. In addition, by including additional behavioural measures (e.g. consumption frequency) future work could investigate the factors that mediate edibility information in the brain, disentangling the roles of appeal and experience.

### 4.5. Conclusion

This EEG study revealed that hunger does not influence the brain’s processing of food-related information, including flavour and personal appeal. Whereas food features such as edibility, food identity, and the flavour profile are encoded relatively early and regardless of whether the food item is relevant, subjective qualities, like current appeal and arousal, only emerge later and are influenced by task relevance. Appeal and arousal reflected personal and dynamic preferences, with limited generalisation across sessions and participants. This study provides insight into the time-course of how the brain processes different aspects of food information, from basic edibility and identity to more complex attributes like flavour and personal appeal.

**Supplementary Figure 1.**
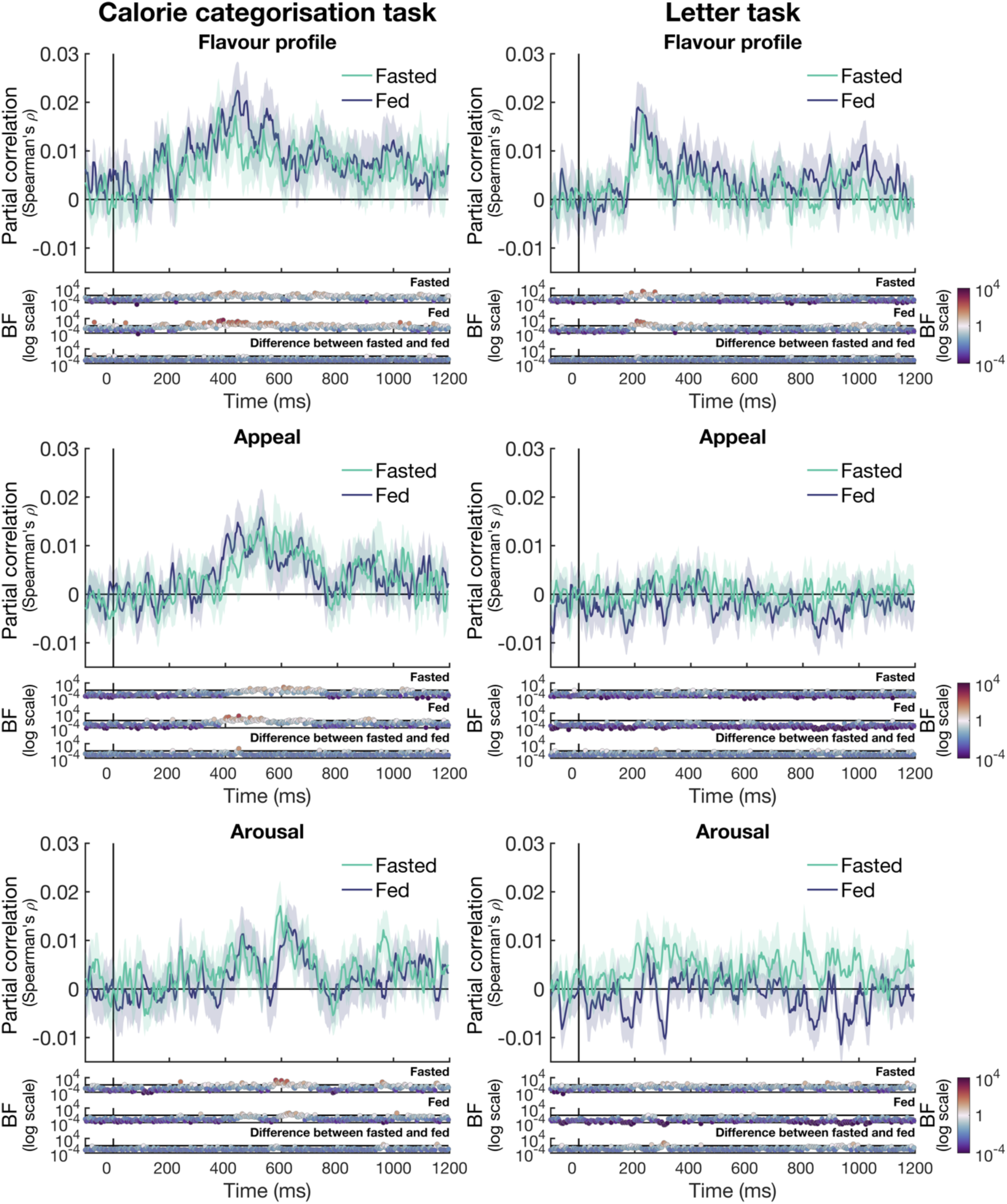
No effect of hunger on the representation of information about the flavour profile, appeal or arousal. We calculated partial correlations between the EEG data for the calorie categorisation task (left) and the letter task (right) and the flavour profile (top), appeal (middle), and arousal (bottom) models. We did this separately for the fasted session (green) and fed session (purple). The shaded areas show the standard error of the mean, plotted in the same colour. We partialled out three CORnet control models from the correlations to control for possible low-level visual differences between the stimuli. The Bayes Factors (BFs) below the plot show evidence for a correlation above 0 in warm colours (BF > 1) and evidence for no correlation in cool colours (BF < 1).

**Supplementary Figure 2.**
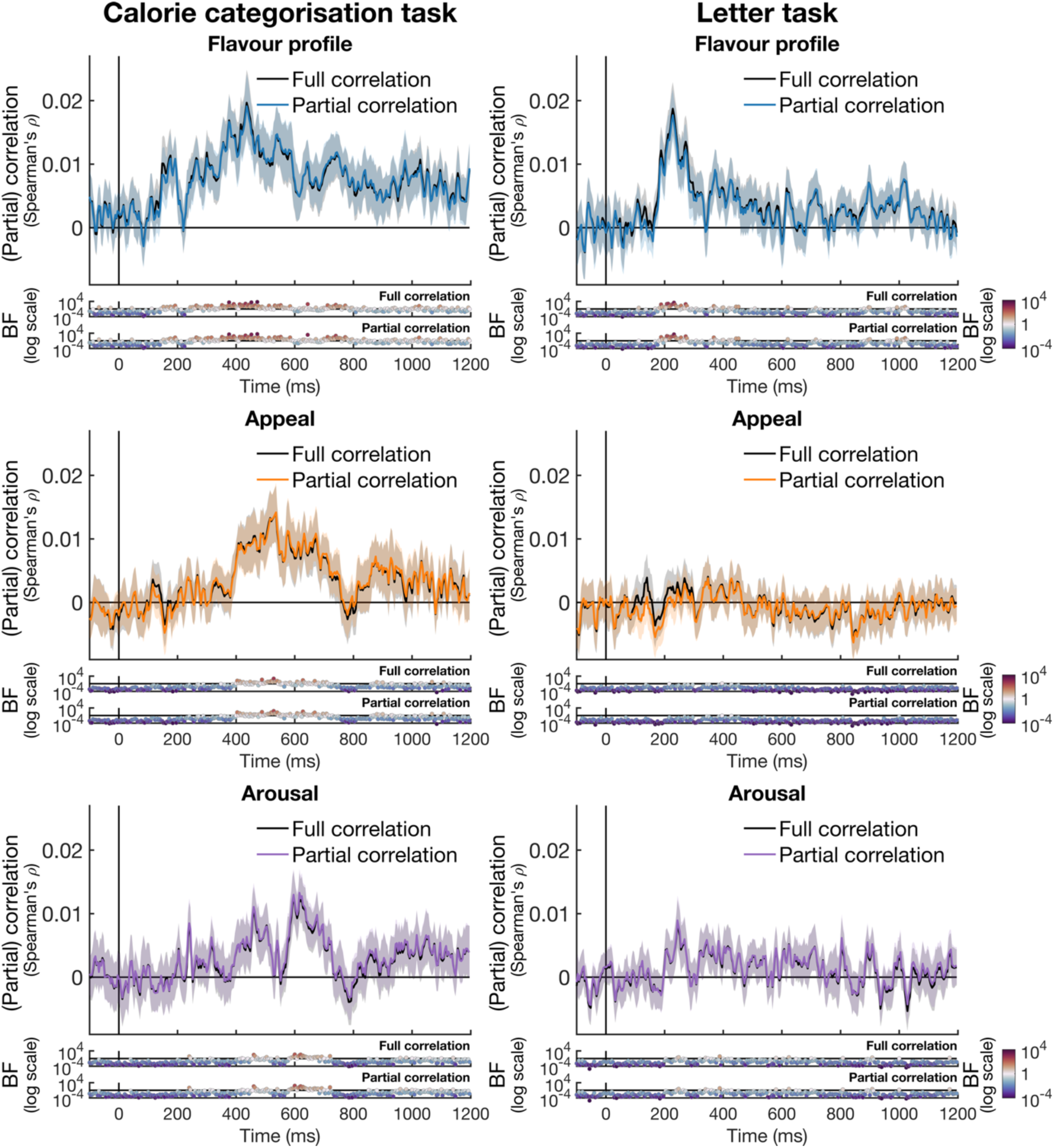
The time-course of full and partial correlations. The full correlations are shown in black. The partial correlations between the EEG data and the models are shown in colour, with the flavour profile shown in blue, the appeal model shown in orange, and the arousal model shown in purple. Left plots show the data for the calorie categorisation task and right plots for the letter task. Bayes Factors for t-tests against a correlation of 0 are shown below each plot, with the top row showing Bayes Factors for the full correlations and the bottom row showing Bayes Factors for the partial correlations.

## Notes

https://doi.org/10.17605/OSF.IO/ZFD7P

### Competing Interest Statement

The authors have declared no competing interest.

### Summary of Updates

We have added further clarifications throughout.

### Funding

- Australian Research Council: DP200101787

